# Recruitment of Two Dyneins to an mRNA-Dependent Bicaudal D Transport Complex

**DOI:** 10.1101/273755

**Authors:** Thomas E. Sladewski, Neil Billington, M. Yusuf Ali, Carol S. Bookwalter, Hailong Lu, Elena B. Krementsova, Trina A. Schroer, Kathleen M. Trybus

## Abstract

We investigated the role of binding partners of full-length *Drosophila* Bicaudal D (BicD) in the activation of dynein-dynactin motility for mRNA transport on microtubules. In single-molecule assays, full-length BicD robustly activated dynein-dynactin only when both the mRNA binding protein Egalitarian (Egl), and *K10* mRNA cargo were present. Electron microscopy showed that both Egl and mRNA were needed to disrupt an auto-inhibited, looped BicD conformation that sterically prevents dynein-dynactin binding. *In vitro* reconstituted messenger ribonucleoprotein (mRNP) complexes with two Egl molecules showed faster speeds and longer run lengths than mRNPs with one Egl, suggesting that cargo binding enhances dynein recruitment. Labeled dynein showed that BicD can recruit two dimeric dyneins to the mRNP, resulting in faster speeds and longer run lengths than with one dynein. The fully reconstituted mRNP provides a model for understanding how adaptor proteins and cargo cooperate to confer optimal transport properties to a dynein-driven transport complex.

## Introduction

Mammalian cytoplasmic dynein-1 (hereafter dynein) is a 12 subunit, 1.4 MDa dimeric molecular motor complex of the AAA+ ATPase family that provides essential cellular functions including transport of vesicles, organelles, and mRNA (reviewed in (Roberts et al., 2013)). Dynein is the predominant minus-end directed microtubule-based motor that traffics cellular cargoes for many microns at speeds of ∼1 μm/sec (Allan, 2011). Molecular motors that transport cargo typically exhibit processive behavior when assayed *in vitro*, meaning that single motors remain bound to the polymer track for long distances without dissociating. It was therefore surprising when *in vitro* studies showed that single molecules of mammalian dynein were at best weakly processive, even in the presence of the multi-subunit 1.2 MDa dynactin complex that is needed for most cellular functions of dynein (King and Schroer, 2000; McKenney et al., 2014; Miura et al., 2010; Ross et al., 2006; Schlager et al., 2014; Trokter et al., 2012; Wang et al., 1995). It was not initially recognized that dynein, like other molecular motors, exists in an auto-inhibited state. One of the first structural studies that investigated the auto-inhibition of dynein showed that the motor domain rings are closely stacked together in a structure called the phi particle (Torisawa et al., 2014). Several non-physiologic mechanisms of disrupting this interaction, such as coupling multiple dyneins to a DNA origami (Torisawa et al., 2014) or binding to a bead (Belyy et al., 2016; King and Schroer, 2000; Mallik et al., 2004), converted dynein from a diffusive to a processive motor by separating the stacked rings.

A major advance in understanding dynein function came from single molecule studies, which showed that binding an α-helical coiled-coil N-terminal fragment of the mammalian adaptor protein Bicaudal-D2 (BicD2^CC1^) to dynein-dynactin caused dynein to become highly processive, with 5-10 μm run lengths and motor accumulation at the microtubule minus-end (McKenney et al., 2014; Schlager et al., 2014). This minimal tripartite complex is called DDB^CC1^ (dynein-dynactin-Bicaudal D2^CC1^). Dynein in the activated DDB^CC1^ complex produced 4.3 pN of force (Belyy et al., 2016), considerably higher than the 0.5-1.5 pN force reported earlier for dynein alone (Mallik et al., 2004; McKenney et al., 2010; Ori-McKenney et al., 2010; Rai et al., 2013), thus allowing it to successfully engage in a tug-of-war with a single kinesin. Recent EM studies provided further insight into the structural basis for the inhibition and activation of dynein. Cryo-EM studies showed that when the dynein ring structure is self-dimerized in the phi particle it locks dynein into a conformation with low affinity for microtubules; disruption of the motor domain dimerization creates an “open” state that is still inhibited for motion ((Zhang et al., 2017). Only when bound to dynactin and an adaptor such as BicD2^CC1^ are the dynein motor domains aligned in a parallel orientation on the microtubule that supports highly processive movement (Chowdhury et al., 2015).

Cargo adaptor proteins such as BicD are thus central to understanding how dynein is activated in the cell. BicD2^CC1^ artificially fused to mitochondria or peroxisome anchoring sequences supports robust dynein-driven motility in HeLa cells. Full-length BicD, in contrast, has only a mild effect on organelle re-localization (Hoogenraad et al., 2003), presumably because it is auto-inhibited and does not bind to or activate dynein-dynactin. BicD2 is composed of three α-helical coiled-coil domains; the N-terminal domain (CC1) is involved with dynein-dynactin binding and activation (Urnavicius et al., 2015), whereas the C-terminal domain (CC3) binds adaptor proteins that link dynein to cargo (Liu et al., 2013). Early yeast-2-hybrid studies showed that the CC1 domain interacts with the CC3 domain, leading to a model in which BicD forms N-to C-terminal auto-inhibitory interactions (Hoogenraad et al., 2001). Early metal-shadowed images of BicD, in contrast, appeared more consistent with a CC2-CC3 interaction (Stuurman et al., 1999). One hypothesis for cellular activation of dynein-dynactin by BicD is that cargo-binding proteins relieve the auto-inhibition of BicD.

Here we test this cargo-activation model by *in vitro* reconstitution of a biologically well-characterized system, namely localization of *K10* mRNA in *Drosophila*. During development, *K10* mRNA is transported by dynein from nurse cells to the *Drosophila* oocyte, where it localizes to the anterior margin to establish the dorso-ventral axis of the *Drosophila* egg (Cheung et al., 1992). *K10* contains a single 44 base-pair transport/localization element (TLS) in its 3’UTR that binds the mRNA-binding adaptor protein Egalitarian (Egl), which in turn binds to the CC3 domain of *Drosophila* BicD (Dienstbier et al., 2009; Serano and Cohen, 1995).

To understand the molecular basis for dynein activation in this system, we reconstituted a mRNP (messenger ribonucleotide protein) complex *in vitro* from purified dynein, dynactin, full-length BicD, Egl, and synthesized *K10* mRNA. Electron microscopy of negatively stained full-length BicD followed by image averaging revealed an auto-inhibited, looped BicD structure which persists in the presence of Egl, but is disrupted once mRNA is bound. Single molecule techniques showed that only mRNA-bound Egl supports the recruitment of dynein-dynactin to full-length BicD, and the consequent movement of the mRNP. Binding of two Egl molecules to BicD favors recruitment of two dynein motors to the DDBE-mRNA complex, and results in faster and longer motion than when only one dynein is bound. These results suggest that multiple interactions contribute to optimal motion of the mRNP on the microtubule.

## Results

### Full-length BicD forms an auto-inhibited looped conformation that does not bind dynein-dynactin

A single molecule pull-down assay was used to determine if full-length *Drosophila* BicD (hereafter called BicD) binds to dynein-dynactin *in vitro*. BicD is composed of predicted α-helical coiled-coil domains shown in blue (CC1, CC2, and CC3) separated by non-coiled coil regions shown in gray (**Figure 1A**). Dynein-dynactin mixtures were incubated with either truncated BicD2^CC1^ that is known to bind and activate dynein-dynactin (McKenney et al., 2014; Schlager et al., 2014), or full-length BicD (**Figure 1B**). Qdot labeled-BicD constructs, along with dynein and dynactin were applied to flow cells with surface-adhered microtubules in the presence of the ATP analog AMP-PNP, which causes dynein to bind strongly to microtubules. Control experiments showed that neither BicD2^CC1^ nor BicD in the absence of dynein-dynactin interacted non-specifically with microtubules. BicD association with microtubules is thus a measure of a tripartite BicD-dynein-dynactin complex. Total internal reflection fluorescence (TIRF) microscopy was used to visualize dynein-dynactin complexes bound to BicD2^CC1^ or BicD (**Figure 1C**). BicD2^CC1^ showed a 13-fold enhanced recruitment to dynein-dynactin bound to microtubules compared with full-length BicD (**Figure 1D**).

**Figure 1.**
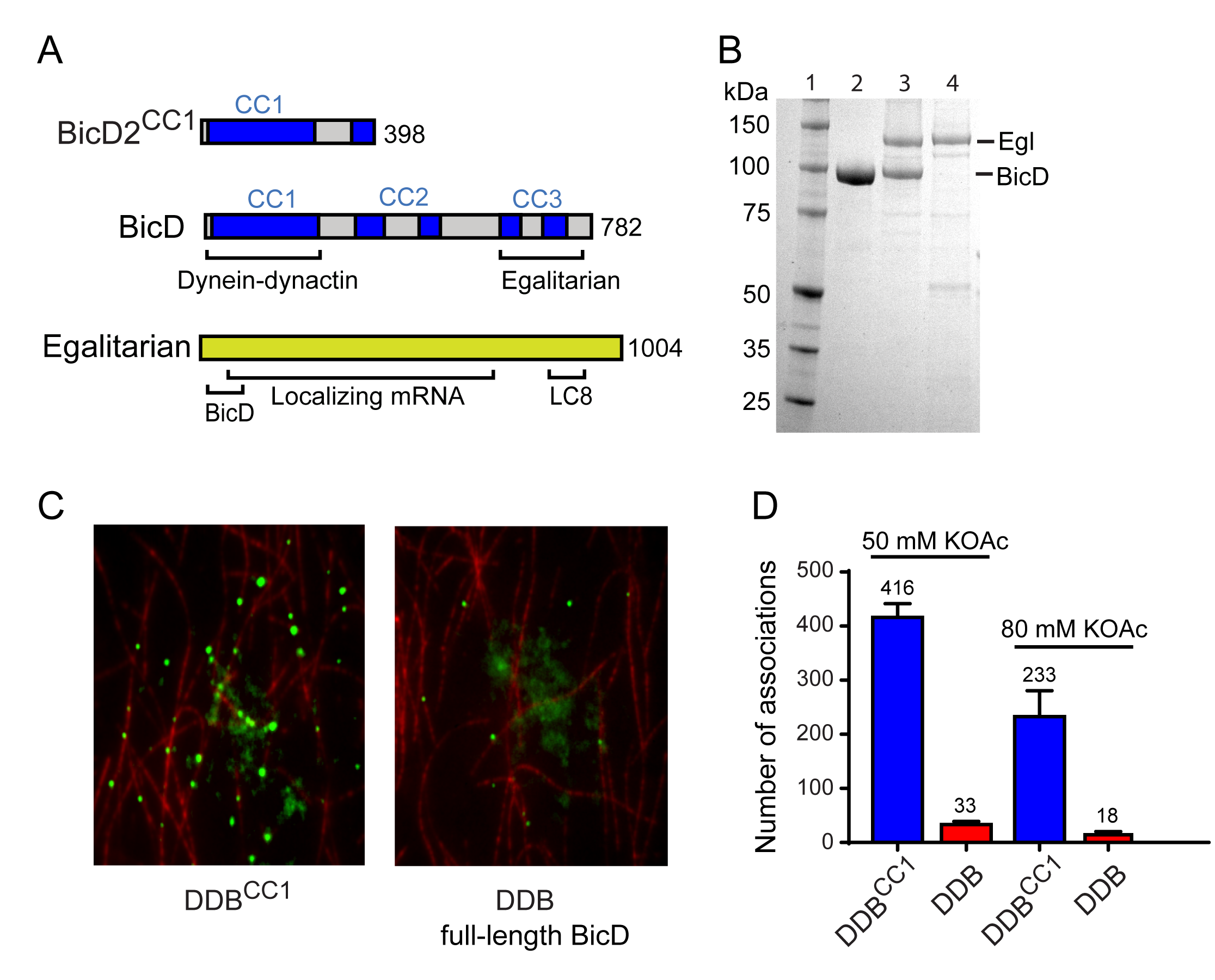
Full-length BicD is auto-inhibited and does not bind dynein-dynactin. (A) Schematic of adaptor proteins. Truncated BicD (BicD2^CC1^) is the minimal fragment that activates dynein-dynactin, but does not bind adaptor proteins. Regions predicted to form coiled-coil (CC) domains from Paircoil2 analysis (McDonnell et al., 2006) are shown in blue. The coiled-coil regions in full-length *Drosophila* BicD are designated CC1, CC2, and CC3. CC2 and CC3 are each interrupted by non-coiled-coil regions (gray). CC1 binds dynein-dynactin and CC3 binds Egalitarian (Egl). Egl has an N-terminal BicD-binding domain, and interacts with dynein light chain (LC8) at its C-terminus. mRNA binding is mediated through a large number (∼800) of amino acids. **(B)** SDS-PAGE gel of (lane 1) molecular mass markers, (lane 2) BicD, (lane 3) BicD co-expressed with Egl, and (lane 4) Egl. 4-12% SDS-gel, MOPS buffer. **(C)** Single molecule pull-downs on microtubules (red) show that dynein-dynactin associates with BicD2^CC1^ (green) but not with full-length BicD (green) in the presence of AMP-PNP. BicD was visualized with a 525 nm-streptavidin Qdot bound to an N-terminal biotin tag on BicD. **(D)** Quantification of the number of dynein-dependent associations of complexes containing BicD2^CC1^ (blue) or full-length BicD (red) to microtubules, normalized to microtubule length and dynein concentration. Data are shown for 50 mM KOAc (left) and 80 mM KOAc (right) ionic strength conditions.

Electron microscopy was used to determine the structural basis for BicD auto-inhibition. Negatively stained EM images of YFP-BicD show two distinct globular densities at the N-terminus that correspond to YFP, confirming the formation of a parallel coiled-coil dimer. The montage (**Figure 2A**) is arranged to show (row 1) the most common “b” shape, (row 2) a less common “d” configuration, (row 3) the range over which the molecule can flex, (row 4) some very compact molecules, and (row 5) rare open molecules. The YFP densities join a fairly straight CC1 region followed by a flexible break in the coiled-coil from which CC2 and CC3 loop back to interact at a point approximately halfway along the CC1 coiled coil (**Figure 2B**). The interpretation of the relatively straight portion of BicD as CC1 is consistent with contour length measurements. The distance from YFP to the very bottom of where the loop begins is 38.7 ± 3.8 nm (±SD, n=150), in good agreement with Paircoil2 analysis (McDonnell et al., 2006) which predicts that the first 258 amino acids of BicD (CC1) have a high propensity for forming a α-helical coiled-coil (0.146 nm rise/residue x 258 residues = 37.7 nm). The contour length from YFP to where the end of the CC2-CC3 loop interacts with CC1 is 16.5 ± 2.3 nm (± SD, n=133), and from this point to the very bottom of the loop is 19.1 ± 2.0 nm (± SD, n=150). The sum of the two approximate halves (16.5 + 19.1 nm) is 35.6 nm, similar to the 38.7 nm measured from YFP to the very bottom of the loop. The contour length of the curved side of the loop is 24.5 ± 2.7 nm (± SD, n=148). These measurements predict that the end of the CC2-CC3 loop interacts with residues near amino acid 113 of BicD. The most common feature of BicD is the loop which appears to be formed by parts of all three coiled coil segments, which is readily seen in averages of all molecules (“global”) as well as averages of the most common “b” type motif molecules (**Figure 2C**). Pull-downs with FLAG antibody resin confirmed that the CC1^HIS^ domain interacts with a construct containing the CC2-CC3^FLAG^ domains, but not with a construct composed only of CC2^FLAG^ (data not shown), consistent with our interpretation of the EM images.

**Figure 2.**
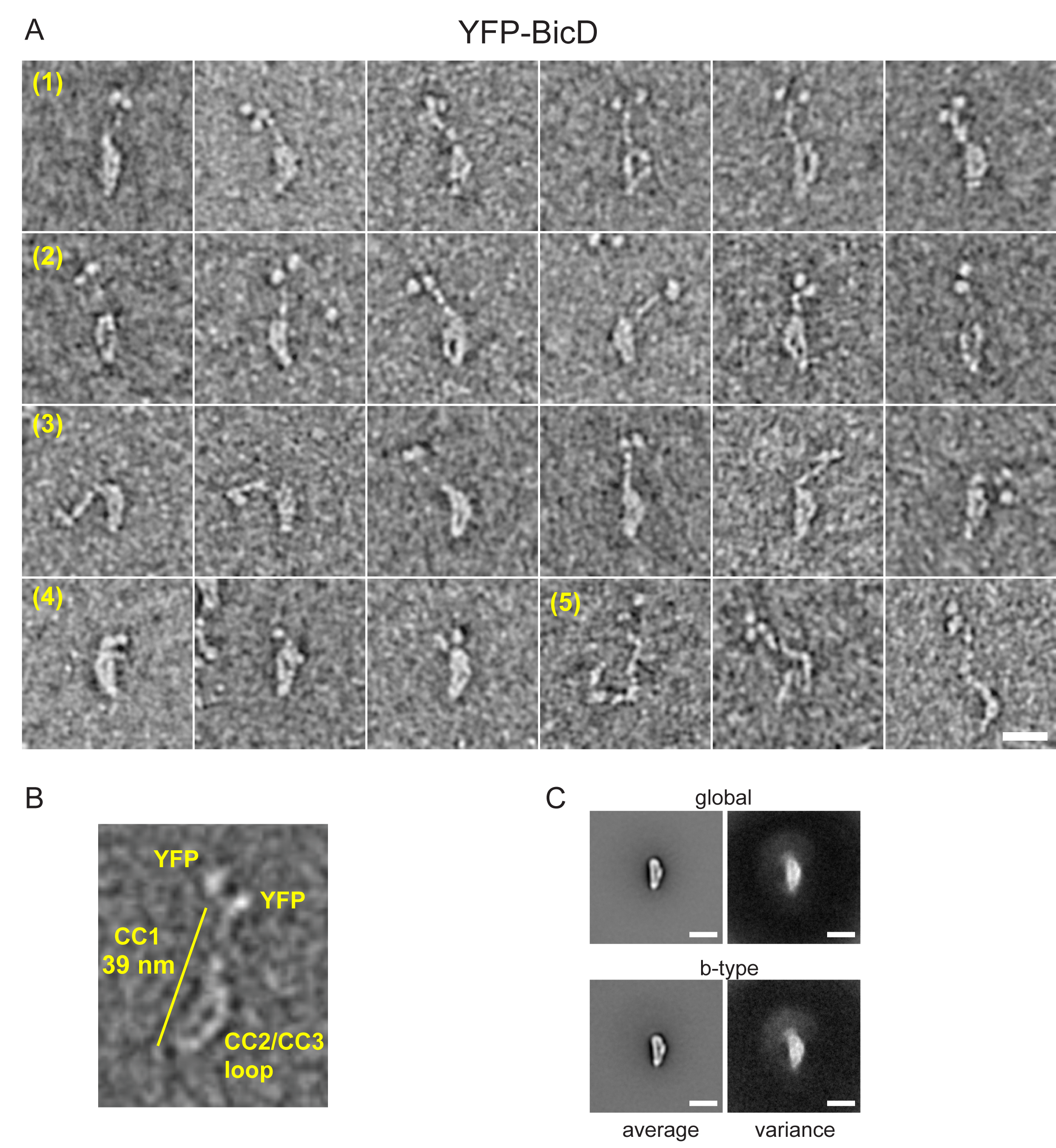
Electron micrographs of the auto-inhibited full-length YFP-BicD. (A) A montage of negatively stained images showing (row 1) the most common “b” shape, (row 2) a less common “d” configuration, (row 3) the range over which the molecule can flex, (row 4) compact molecules, and (row 5) rare open molecules. **(B)** One YFP-BicD molecule illustrating the interpretation of the EM image, with the length of the long straight section indicated. **(C)** Average and variance of all molecules (global), or of only the “b” type configuration molecules, both of which highlight the common looped part of the molecule. Scale bars=20 nm.

### Binding of Egl does not disrupt the auto-inhibited structure of BicD

BicD and the mRNA binding protein Egl can either be expressed separately and then reconstituted after purification, or co-expressed as a complex in Sf9 cells (**Figure 1B**). Electron microscopy of the co-expressed BicD-Egl complex revealed a remarkably similar overall conformation to that of BicD alone, with the loop conformation remaining intact, and the two N-terminal YFP globular domains on BicD confirming the presence of a parallel coiled-coil. An additional globular structure was often observed adjacent to the loop (**Figure 3A**). The N-terminal region of Egl (amino acids 1-79) binds BicD, and residues 557-726 of Egl contain a globular exonuclease homology region (Dienstbier et al., 2009), which may be the globular structure visualized here. Averages of the “b” type motif molecules (**Figure 3B**) are very similar to those seen in the absence of Egl. Co-alignment of the BicD-Egl images with images of BicD alone revealed that the major difference in density is located in a region towards the bottom of the loop, suggesting that this region is an attachment site for Egl (**Figure 3C**). Some raw images show an attachment to the top of the loop but this was not seen in the subtraction of the aligned images. The heatmap (**Figure 3C**) suggests that the extra globular domain of Egl can occupy a range of positions relative to the BicD loop, consistent with single particle images. The lack of a distinct density in the difference map, and the distal and highly variable location of the additional density seen in images of BicD-Egl particles suggest the existence of a long flexible linker between the globular domain and the N-terminal region of Egl that binds BicD. **Video 1** shows a low resolution 3D map of negative stain EM data, which allows size comparison of the loop with existing structures for parts of the YFP-BicD-Egl complex.

**Figure 3.**
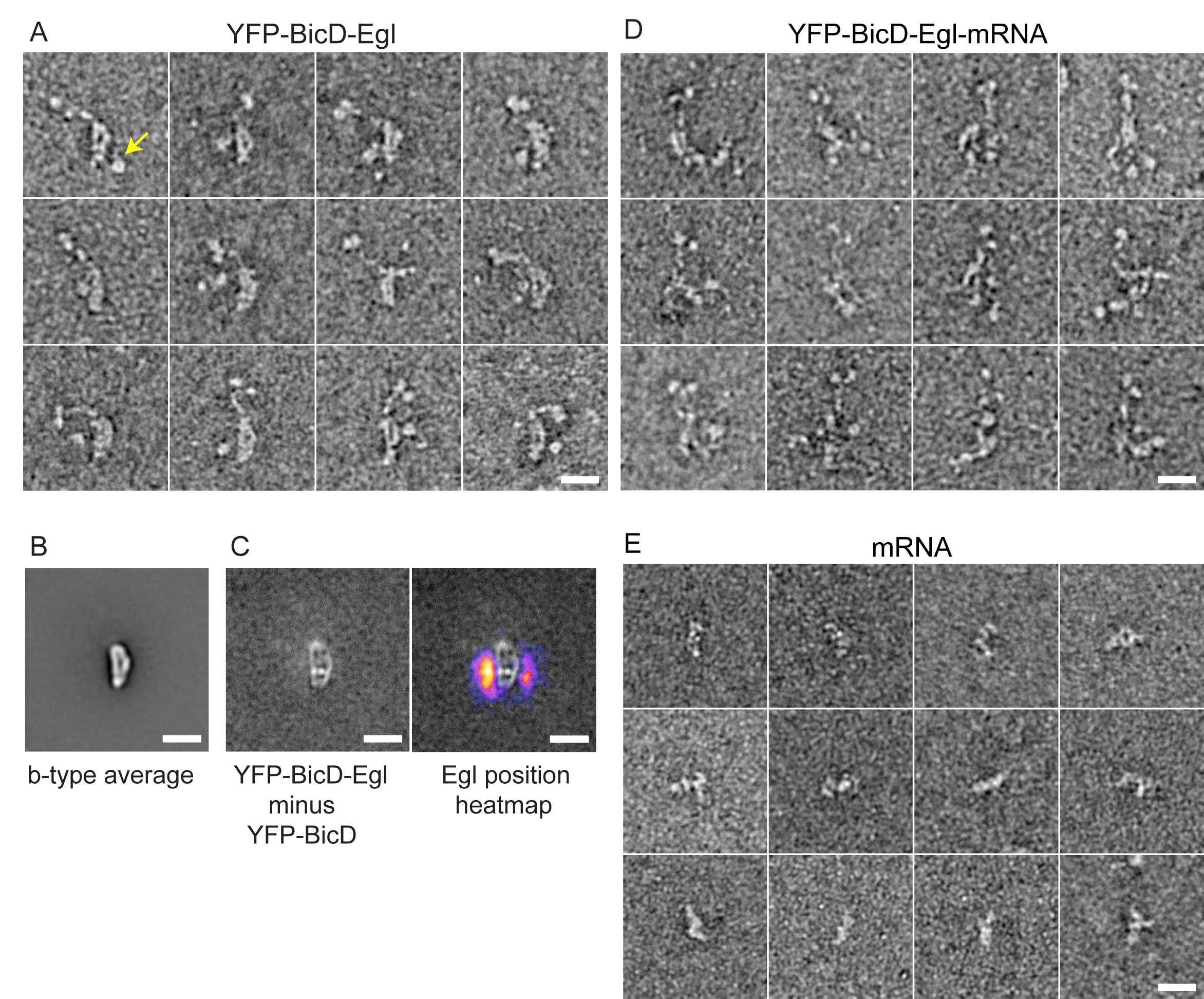
Electron micrographs of the YFP-BicD-Egl and YFP-BicD-Egl-mRNA complex. (A) A montage of negatively stained images showing that BicD retains the auto-inhibited looped conformation in the presence of bound Egl. **(B)** Average of all YFP-BicD-Egl “b” type molecules. **(C)** (Left image), a subtraction image showing the average of all “b” type images of YFP-BicD-Egl minus the average of all “b” type images of YFP-BicD alone. (Right image), the same image with a heatmap of the position of the globular Egl domain overlaid. **(D)** A montage of negatively stained images showing that BicD no longer retains the auto-inhibited looped conformation in the presence of bound Egl and mRNA (*K10*min). **(E)** A montage of negatively stained images of the mRNA (*K10* min) alone. Scale bars=20 nm.

The EM images imply that Egl alone is not sufficient to disrupt the auto-inhibited conformation of BicD. This result predicts that the BicD-Egl complex will not bind to dynein-dynactin. Single molecule pull-downs confirmed that full-length YFP-BicD and Egl-Qdot were not recruited to microtubule-bound dynein-dynactin (**Figure 4A, B**).

**Figure 4.**
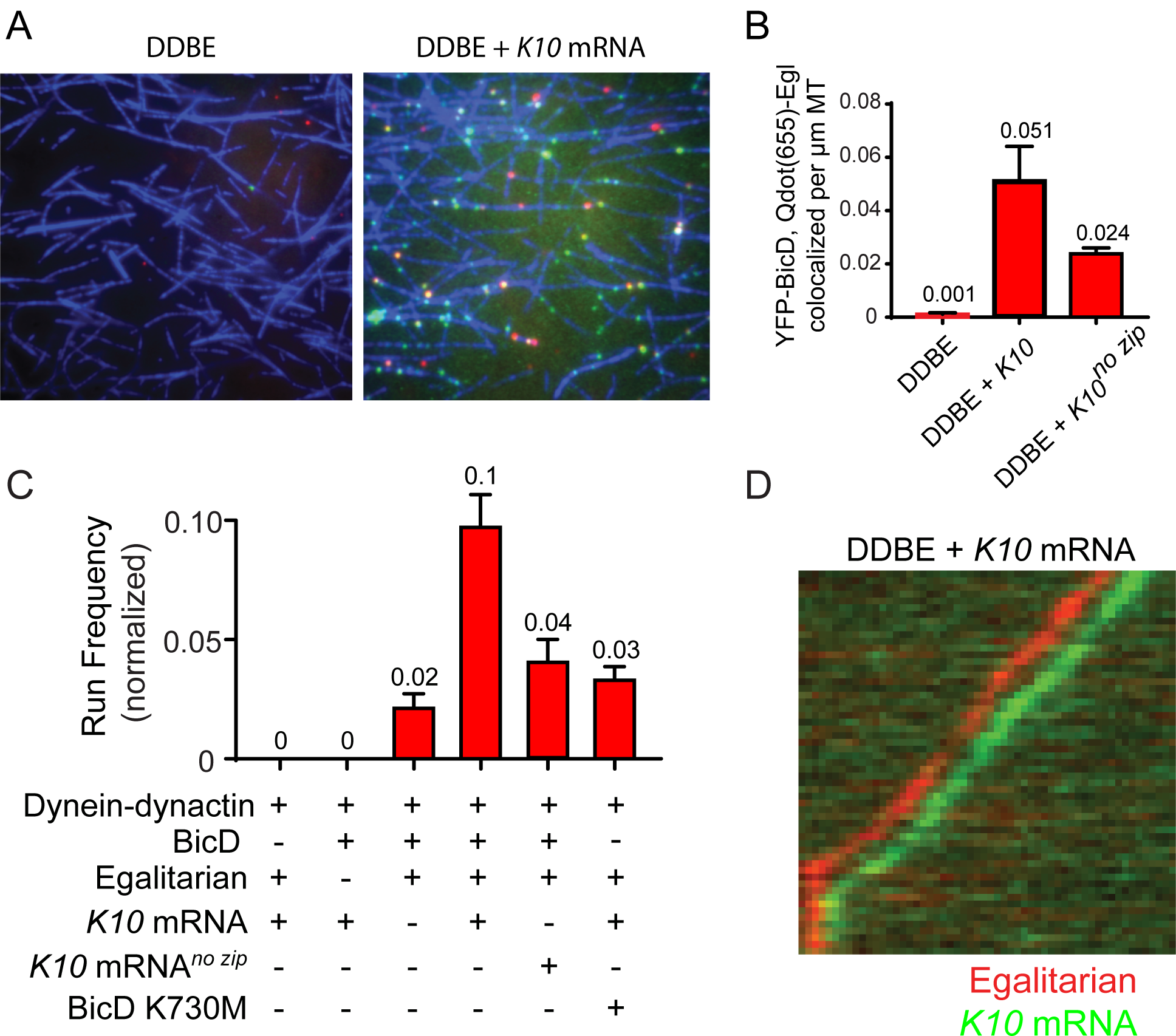
*K10* mRNA is needed for BicD-Egl to recruit dynein-dynactin for motility. (A) Single molecule pull-downs of YFP-BicD (green) and Qdot-Egl (red) by dynein-dynactin (DDBE) bound to microtubules (blue) in the absence or presence of *K10* mRNA. **(B)** Quantification of the single molecule pull-down data showing the number of co-localized BicD-Egl complexes bound to dynein-dynactin (normalized per μm microtubule length) in the absence of mRNA (DDBE), in the presence of *K10* mRNA (DDBE + *K10*), or *K10* without the TLS zip code (DDBE + *K10*^no zip^). **(C)** Run frequencies (n runs per μM dynein per μm microtubule length per s) of motile mRNPs. (Bars, left to right) Movement of labeled *K10* mRNA is not observed in the absence of either BicD or Egl. Few events are observed for mRNPs lacking mRNA (movement was visualized with a Qdot bound to Egl). Fully reconstituted mRNPs with *K10* mRNA have the highest run frequency, while mRNPs reconstituted with *K10* mRNA lacking the TLS zip code showed reduced run frequencies. mRNPs reconstituted with a mutant of BicD (BicD K730M) that has impaired binding to Egl shows fewer events than WT Egl. Error bars, sem, n = 4 movies per condition. Data are representative of 3 independent preparations of dynein-dynactin. **(D)** Kymograph showing DDBE and K10 mRNA dual labeled with Egl bound to a Qdot (red) and *K10* mRNA labeled with Alexa Fluor 488 UTP (green). Color channels are offset for presentation purposes.

### The BicD-Egl complex binds and robustly activates dynein-dynactin in the presence of mRNA

We next tested whether an mRNA cargo was required for BicD-Egl to bind to dynein-dynactin, by reconstituting an mRNP consisting of dynein-dynactin, BicD, Egl and a *K10* mRNA transcript. The presence of *K10* mRNA enhanced Egl and BicD co-localization with microtubule-bound dynein-dynactin (**Figure 4A, B**). This observation implies that both mRNA and Egl are needed for BicD to adopt a conformation that can recruit dynein-dynactin. Consistent with this interpretation, electron microscopy of the BicD-Egl-*K10*_min_ mRNA complex showed a variety of flexible structures, none of which retained the inhibitory looped conformation seen for BicD alone or the BicD-Egl complex (**Figure 3D versus Figure 2A, 3A; Fig. 3E** shows *K10*_min_ mRNA alone). The highly variable appearance of the BicD-Egl-mRNA complex precluded image averaging or detailed assignment of the position of each of the components within the complex.

Single-molecule pulldowns were further used to assess the requirement for a zip code in *K10* mRNA for dynein-dynactin recruitment by BicD-Egl-mRNA complexes. mRNPs reconstituted with a mutant *K10* mRNA transcript lacking the transport/localization sequence (TLS) caused a ∼2-fold reduction in the number of BicD-Egl complexes associated with microtubule-bound dynein-dynactin, compared with native *K10* (**Figure 4B**). Egl thus binds primarily to the TLS zip code, although it is able to bind weakly to other regions of the mRNA.

The run frequencies of reconstituted complexes were determined by TIRF microscopy using either Qdot-labeled adaptor proteins (BicD or Egl), or *K10* mRNA synthesized with Alexa Fluor 488-UTP. In the absence of either BicD or Egl, no movement of labeled *K10* mRNA was observed (**Figure 4C**). In the presence of both BicD and Egl (labeled adaptors), but in the absence of mRNA, only a low frequency of runs was observed. Addition of *K10* mRNA to the dynein-dynactin-BicD-Egl complex enhanced run frequency 5-fold. Consistent with single molecule pull-downs, *K10* mRNA constructs lacking the TLS showed a reduced run frequency compared with wild type *K10* transcripts. In the cell, robust localization of *K10* mRNA requires the TLS zip code (Bullock and Ish-Horowicz, 2001; Serano and Cohen, 1995). Lastly, a mutant of BicD (K730M) which has reduced association with Egl (Dienstbier et al., 2009) supports motility with a reduced frequency compared with wild type BicD. As an additional control, we confirmed that motile *K10* mRNA complexes were bound to Egl by simultaneously tracking labeled *K10* mRNA and Qdot-labeled Egl (**Figure 4D**). Consistent with our data showing that Egl is needed for mRNP motility, *K10* mRNA and Egl co-localize in motile mRNP complexes on microtubules.

### *K10* mRNA activates mRNPs for processive movement with fast speeds

Both the constitutively activate truncated DDB^CC1^ complex and the fully reconstituted mRNP complex (dynein-dynactin-full length BicD-Egl-*K10* mRNA) show long processive runs and accumulation at the minus-end of microtubules (**Figure 5A;** note the vertical lines in the kymographs corresponding to microtubule minus ends). Speeds and run lengths of motile complexes were tracked with high temporal (0.2 s) resolution. *K10* mRNP motility speed (0.45 ± 0.21 μm/sec, n=1126) was faster than the minimal DDB^CC1^ complex (0.35 ± 0.19 μm/sec, n=1147, p<0.05, t-test, mean ± SD) (**Figure 5B**). Run lengths of the *K10* mRNP (7.2 ± 0.5 μm, n=142) were also longer than the minimal DDB^CC1^ complex (5.4 ± 0.7 μm, n=137; p=0.053, Kolmogorov–Smirnov test, mean ± SE) (**Figure 5C**). The fully reconstituted mRNP is thus at least as active as the minimal DDB^CC1^ complex.

**Figure 5.**
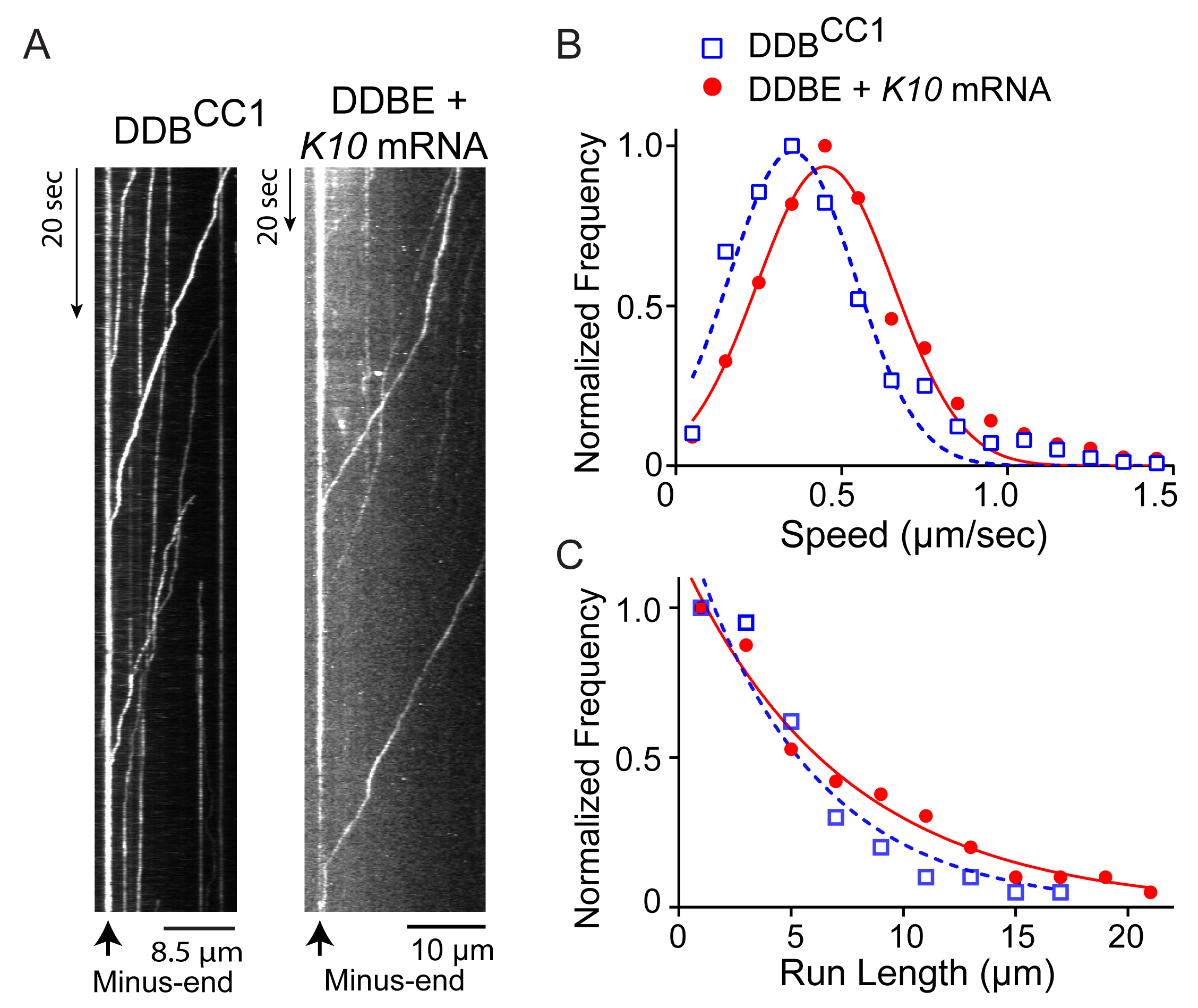
The motile properties of the fully reconstituted mRNP are similar to DDB^CC1^. (A)Kymograph (time vs displacement) of (left panel) a minimal dynein-dynactin-BicD2^CC1^ (DDB^CC1^) complex, visualized with a Qdot bound to BicD2^CC1^, and (right panel), a complex of dynein, dynactin, full-length BicD, Egl and *K10* mRNA labeled with Alexa Fluor 488 UTP. The straight vertical line at the left of both kymographs shows complex accumulation at the minus-end of the microtubule. **(B)** Speed distributions of DDBE + *K10* mRNA (red circles)(0.45 ± 0.21 μm/sec, n = 1126) and DDB^CC1^ (blue squares)(0.35 ± 0.19 μm/sec, n = 1147; p< 0.05, t-test, from 50 representative run trajectories for each condition, mean ± SD). **(C)** Run length distributions of DDBE + *K10* mRNA (red circles) (7.2 ± 0.5 μm, n=142) and DDB^CC1^ (blue squares)(5.4 ± 0.7 μm, n = 137) (p=0.053, Kolmogorov–Smirnov test, mean ± SE of the fit).

### Role of LC8

We tested if the proposed interaction of Egl with LC8 on dynein affects the function of the mRNP. Egalitarian binds to the dynein light chain LC8 (DDLC1 in *Drosophila*) through a consensus light chain binding site on Egl between amino acids 963-969 (^963^AESQTLS^969^)(Navarro et al., 2004). The S965L mutation inhibits light chain binding but does not interfere with either its ability to bind mRNA or BicD, yet it compromises mRNA transport in the cell (Dienstbier et al., 2009; Navarro et al., 2004). Our data show no statistical difference in speed between complexes reconstituted with WT versus mutant S965L Egalitarian (0.43 ± 0.53 μm/sec, n=850 for Egl^WT^, versus 0.47 ± 0.15 μm/sec, n= 641 for the mutant Egl^mut^; p*=*0.12, t-test, mean ± SD). Likewise, there was no statistical difference between the run lengths (6.2 ± 0.211 μm, n=74 for Egl^WT^ versus 5.0 ± 0.36 μm for Egl^mut^, n=70, p=0.6, Kolmogorov–Smirnov test, mean ± SE).

Because we could not prove that the point mutation actually affected the potential interaction between LC8 and Egl in the context of the entire mRNP, we also expressed dynein without LC8, and reconstituted mRNPs containing it. Our data showed no statistical difference in speed between mRNPs reconstituted with dynein, versus dynein minus the LC8 light chain (0.41 ± 0.13 μm/sec, n=396 for dynein, versus 0.42 ± 0.13 μm/sec, n= 488 for dynein without LC8; p*=*0.46, t-test, mean ± SD). Thus the putative interaction between Egl and the LC8 light chain of dynein has no obvious functional effect.

### Motile mRNPs contain multiple copies of Egl

BicD is a coiled-coil, and thus has the potential to bind two molecules of Egl. To test whether the speed and run length of the mRNP depends on Egl stoichiometry, Egl with a C-terminal biotin tag was separately labeled with either a green (565 nm) or a red (655 nm) streptavidin-Qdot, then blocked with excess biotin to prevent further binding. *K10* mRNPs were then reconstituted with equimolar red and green labeled Egl (**Figure 6A**). Motile complexes containing both green and red Qdots were observed, as were motile complexes with a single color (**Figure 6B**). The complexes with only one color are likely a mixture of complexes containing only one Egl, or two Egls of the same color. In rare cases, a moving complex started dual-colored, and then reduced to a single color in the same trajectory (data not shown), indicating that one Egl is sufficient to support a motile mRNP.

**Figure 6.**
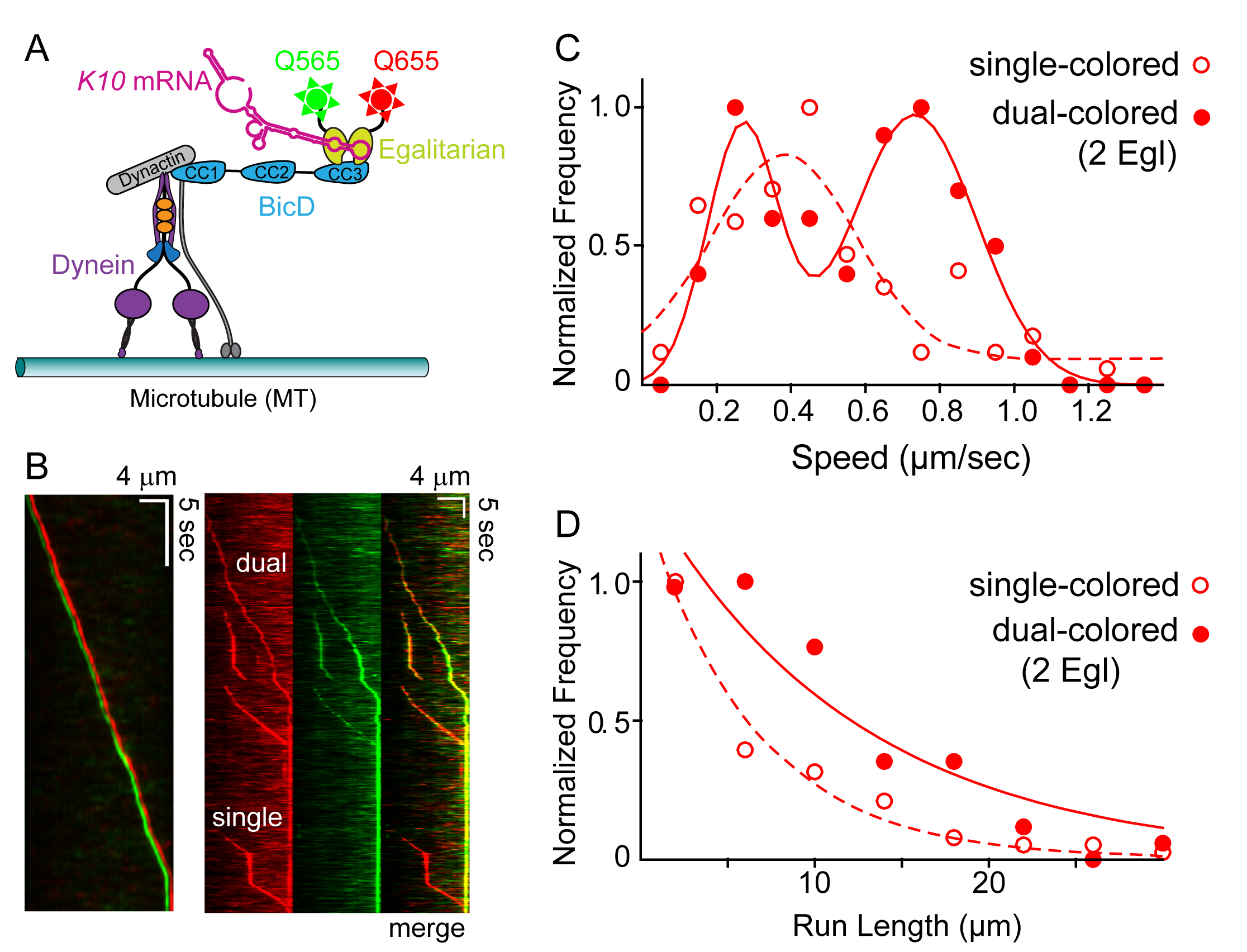
Motile properties of complexes containing one versus two Egl molecules. **(A)** Schematic of the two-color experiment in which Egl is labeled with either a 565 or 655 nm QDot. **(B)** Kymographs of dual color runs. (Left panel), Trajectories of a run with both a 565 nm(green) and a 655 nm (red) Qdot-labeled Egl bound to the moving complex. The red trajectory is shifted horizontally for presentation purposes. (Right panels) Red, green and combined channels of several processive runs. The top 3 trajectories are dual color runs, while the bottom one is a single color run. **(C)** Speeds of single color complexes (open red circles) were best fit to a single Gaussian distribution with a speed of 0.46 ± 0.07 μm/sec (n= 81). In contrast, the dual colored complex (filled red circles) was best fit with a double Gaussian distribution with a slow speed of 0.27 ± 0.1 μm/sec, and a faster speed of 0.73 ± 0.16 μm/sec (n= 62). **(D)** Run length histogram of single color events (open red circles)(6.4 ± 0.7 μm, n=81) are significantly shorter than dual color runs (filled red circles) (12.1 ± 2.4 μm, n=62; p=0.045, Kolmogorov– Smirnov test, mean ± SE of the fit). The curves are exponential fits to the data.

The speed and run length distributions of the single versus dual-colored complexes differed. The speed histogram of the dual colored complex was best fit with a double Gaussian distribution with a slow speed of 0.27 ± 0.1 μm/sec, and a faster speed of 0.73 ± 0.16 μm/sec (n= 62). Speeds of the single colored complexes were better fit to a broad single Gaussian distribution, with a speed of 0.46 ± 0.07 μm/sec (n= 81), despite the fact that some single colored events may contain two Egl molecules of the same color (**Figure 6C**). Run length distributions of the single-colored runs (6.4 ± 0.7 μm, n=81) were significantly shorter than the dual-colored runs (12.1 ± 2.4 μm, n=62; p=0.045, Kolmogorov–Smirnov test) (**Figure 6D**). An explanation that would account for both the enhanced speed and run length is that binding of two Egl molecules favors recruitment of two dimeric dynein motors to the dynactin-BicD-mRNA complex (Grotjahn et al., 2018; Urnavicius et al., 2018). This result is particularly striking in light of the fact that experiments performed up to this point used a molar ratio of ∼1 dynein per dynactin to assembly the mRNP, as this was the assumed stoichiometry of binding.

### mRNPs containing two dyneins move faster and longer

The experiments with dual-colored Egl molecules suggested that BicD has the potential to recruit two dynein motors. To directly test this idea, expressed dynein with an N-terminal biotin tag was separately labeled with either Alexa Fluor 647 (red) or Alexa Fluor 488 (green)(**Figure 7A**). *K10* mRNPs were then reconstituted with equimolar red and green labeled dyneins, with a molar ratio of 2 dyneins per dynactin. Single molecule pulldowns in the presence of AMP-PNP showed 22% dual-colored complexes (n=274), direct evidence that two dynein motors can be recruited (**Figure 7B**). A comparable experiment using dynein-dynactin and truncated BicD2^CC1^ (n=114) showed 23% dual-colored complexes. Similarly, we obtained 25% (n=185) dual-colored dynein complexes with a tripartite complex formed from dynein-dynactin and the adaptor protein Hook1. Hook was used as a control because two recent structural studies agreed that Hook predominantly recruited two dyneins (Grotjahn et al., 2018; Urnavicius et al., 2018). Note that the maximum number of dual-colored complexes that can be statistically obtained with our experimental protocol is 50%.

**Figure 7.**
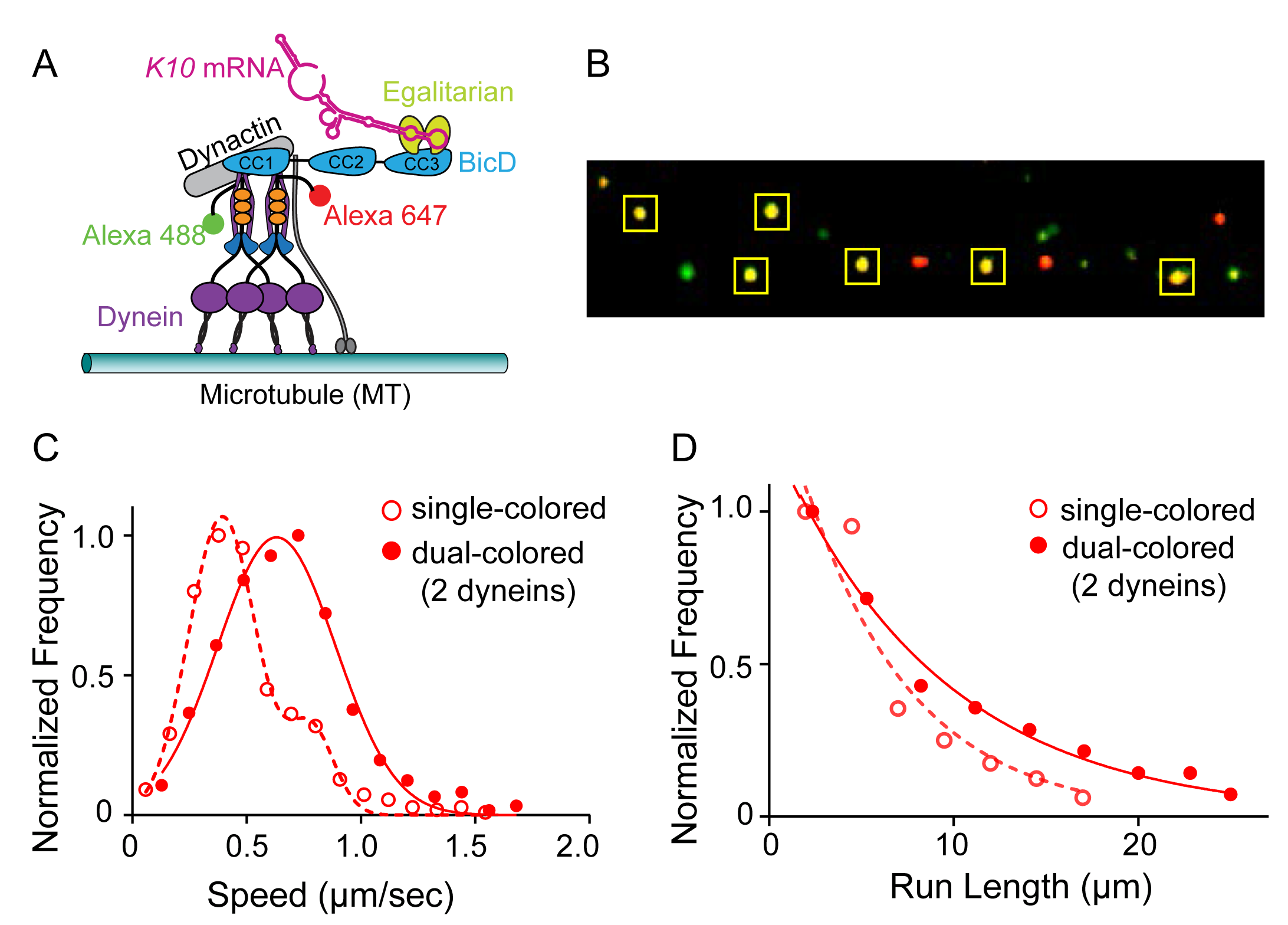
Recruitment of two dynein motors to the mRNP results in faster and longer runs. (A) Schematic of the two-color experiment. Dynein was either labeled with Alexa 488 (green) or Alexa 647 (red) for the single molecule pulldowns shown in panel B. **(B)** Single molecule pulldowns in the presence of AMP-PNP showed that 22% of complexes were dual-colored (maximum of 50%), showing that two dynein motors were bound. **(C)** For the speed and run length data shown in panels C and D, dynein was labeled with either a 525 or a 655nm Qdot. The speed of dual-colored complexes (filled red circles) (0.63 ± 26 μm/sec, n= 40), were compared to that of single-colored complexes (open red circles)(0.40 ± 0.14 μm/sec and 0.72 ± 0.13 μm/sec, n= 44). **(D)** Run lengths of dual-colored runs (filled red circles)(8.9 ± 0.5 μm, n=40) were 53% longer than the single-colored complexes (open red circles)(5.8 ± 1.1 μm, n=44, p=0.036, Kolmogorov–Smirnov test, mean ± SE of the fit).

The motile properties of dual-colored mRNPs that contained two Qdot labelled dyneins, were compared with that of single-colored complexes, which contain either one or two dynein motors. **Video 2** shows an example of a dual-colored complex that moves both faster and longer than single-colored complexes. The speed histogram of the dual-colored complexes was best fit with a single Gaussian distribution with a speed of 0.63 ± 26 μm/sec (n= 40). The single colored complexes were best fit to a double Gaussian distribution with speeds of 0.40 ± 0.14 μm/sec and 0.74 ± 0.13 μm/sec (n= 44), with the faster speed likely corresponding to a smaller population containing two dyneins labeled with the same color (**Figure 7C**). The speed of the dual-colored complexes was 58% faster than the slow speed of the single-colored complex. Run lengths of dual-colored runs (8.9 ±0.5 μm, n=40) were 53% longer than the single-colored complexes (5.8 ± 1.1 μm, n=44, p=0.036, Kolmogorov–Smirnov test) (**Figure 7D**). Recruitment of two dyneins thus results in an mRNP that moves both faster and longer than an mRNP containing only one dynein.

## Discussion

Using purified components, we reconstituted an mRNP composed of dynein-dynactin, BicD, Egl, and a localizing mRNA found in *Drosophila* (*K10*). The complex has the capacity to recruit two dimeric dyneins, and two Egl molecules. Binding of two Egl molecules favors recruitment of two dimeric dyneins, and complexes with two dyneins move faster and longer than those with only one dynein. In the absence of mRNA, full-length BicD, or BicD-Egl, does not bind to dynein-dynactin due to BicD auto-inhibitory interactions. This is in contrast to truncated BicD2^CC1^ that recruits dynein-dynactin and converts it into a highly processive motor with enhanced force output, but which cannot bind cargo (Belyy et al., 2016; McKenney et al., 2014; Schlager et al., 2014). Class averages of negatively stained full-length BicD revealed the structural basis for BicD auto-inhibition: BicD forms a looped conformation that resembles the letter “b”. The long straight portion of the letter “b” is attributed to the first coiled-coil region, CC1 (∼40 nm), and the remaining loop to CC2-CC3. An atomic resolution structure of dynactin in complex with BicD2^CC1^ and the dynein tail showed that 275 amino acids (∼40 nm) of BicD bind along dynactin (Urnavicius et al., 2015). Although essentially all of the ∼40 nm CC1 is exposed in full-length auto-inhibited BicD, binding of CC3 to the middle of CC1 must be sufficient to block formation of a stable high-affinity ternary complex with dynein-dynactin. In this system, both *K10* mRNA and Egl are necessary to disrupt the auto-inhibitory loop and recruit dynein-dynactin for long processive runs on microtubules. The requirement for mRNA ensures that motor activation is coupled to cargo binding, preventing futile dynein activity. A concurrent study from the Bullock laboratory (McClintock et al., 2018) also showed the requirement for mRNA to stabilize a dynein-dynactin-BicD-Egl complex that is capable of supporting high frequency, long processive runs on microtubules.

### Common features of dynein-dynactin adaptor proteins

As more adaptor proteins are studied, both common and unique features begin to emerge. One common feature of known dynein-dynactin adaptors is that they contain an α-helical coiled-coil. Studies on Spindly, a dynein-dynactin adaptor protein that enables kinetochore attachment via a three-subunit complex called Rod-Zw10-Zwilch (RZZ), showed the importance of certain sequences in the Spindly N-terminal region for dynein-dynactin recruitment (Gama et al., 2017; Mosalaganti et al., 2017). These regions are conserved to various extents in adaptors such as BicD2, BicDR1, Hook3 and others. The “CC1 box” near the N-terminus of most adaptors, has been implicated in interacting with the light intermediate chain (LIC) of dynein. Depending on the adaptor, ∼250 amino acids after the CC1 box is the “Spindly motif”, a region that interacts with the pointed end complex (consisting of Arp11, p62, p25, p27) of dynactin. A region in Hook3 containing a Spindly-like motif was shown to be required for *in vitro* activation of a dynein-dynactin-Hook3 complex (Schroeder and Vale, 2016). Thus a picture of how the ternary complex (dynein-dynactin-adaptor protein) is stabilized is beginning to emerge. One feature that appears to be unique to BicD is its auto-inhibition. Full-length Spindly (McKenney et al., 2014) and full-length Hook1 and Hook3 (Olenick et al., 2016) can bind and activate dynein-dynactin without truncation, implying that they are not auto-inhibited. For BicD, auto-inhibition provides a mechanism to couple the presence of cargo mRNA to dynein activation.

### Why is mRNA needed to relieve BicD auto-inhibition?

Our EM images of BicD-Egl suggest that part of the Egl molecule folds into a globular domain which is attached via a linker to the BicD binding site. A candidate for this globular domain is the putative 3’-5’ exonuclease domain identified within Egl (Mach and Lehmann, 1997; Moser et al., 1997; Navarro et al., 2004). Individual residues typically associated with exonuclease activity were shown to be unnecessary for the function of Egl, whereas deletion of the entire domain strongly reduced RNA binding activity. The globular domain may represent an RNA binding module that does not require exonuclease activity for normal function.

How does mRNA binding to Egl disrupt the auto-inhibitory looped structure of BicD? Steric/competitive or allosteric mechanisms can be considered. A competitive mechanism suggests that Egl-mRNA, but not Egl alone, competes effectively with CC1 for a common binding site on CC3. A large portion of Egl is involved in interactions with mRNA, some of which overlap with its N-terminal binding site for BicD CC3 (see **Figure 1A**). A competitive mechanism appears to be the case with Rab6^GTP^, an adaptor protein that links dynein-dynactin-BicD to vesicular cargo. The Rab6^GTP^ and CC1 binding sites on CC3 overlap (Terawaki et al., 2015), implying that both cannot bind simultaneously. Rab6^GTP^ (but not Rab6^GDP^) in solution or bound to artificial liposomes released BicD2 from an auto-inhibited state to promote processive dynein-dynactin motion (Huynh and Vale, 2017). The bound GTP in essence signals to BicD that the Rab is bound to cargo (reviewed in (Grosshans et al., 2006)). In the case of Egl, the mRNA cargo needs to be bound to relieve BicD auto-inhibition.

Alternatively, an allosteric mechanism would postulate that mRNA binding to Egl causes a propagated change in the BicD coiled-coil that weakens the CC3-CC1 interaction (Liu et al., 2013). This possibility is due to an unusual feature of part of the *Drosophila* BicD CC3 structure, in which the same residues in the two chains of the coiled-coil adopt different heptad registries, referred to as “heterotypic” interactions. It was proposed that this region may act as a molecular switch to promote weakening of the CC3-CC1 interaction following cargo binding to the adjacent “homotypic” segment of coiled-coil (Liu et al., 2013).

We previously showed that in an actomyosin-based mRNA transport system in budding yeast, mRNA was also required for processive motion, but for a different reason: mRNA stabilized the interaction of two single-headed class V myosins (Myo4/She3) with the mRNA binding adaptor protein She2 at physiological ionic strength (Sladewski et al., 2013). Although mechanistically the role of mRNA differs from what we show here with dynein-dynactin and BicD, in both cases the requirement for mRNA ensures that only cargo-bound motor complexes are motile.

### Role of the zip code

The highest frequency of moving complexes, and the highest number of complex associations detected in single molecule pull-downs, were obtained with *K10* mRNA containing the TLS zip code. These values decreased ∼two-fold when the zip code was removed, but remained higher than in the absence of *K10* mRNA, implying that Egl may also bind to sequences outside the TLS. This interaction is specific to the *K10* mRNA transcript because the tRNA used as a non-specific blocking reagent in single molecule reconstitutions is not capable of activating DDBE complexes. Studies using extracts from *Drosophila* embryos also concluded that the role of localization signals/zip codes is to increase the average copy number of dynein-dynactin recruited to an mRNP (Amrute-Nayak and Bullock, 2012). The function of zip codes in *Drosophila* appears to differ from that seen in budding yeast, where zip codes in the model mRNA *ASH1* are essential for recruitment of myosin motors to the mRNA binding adaptor protein She2 (Sladewski et al., 2013).

### mRNPs recruit two Egalitarians and two dyneins

Previous studies showed that two Rab6^GTP^ adaptors can bind to the CC3 domain of BicD, by associating with opposite faces of the BicD coiled-coil (Liu et al., 2013). The binding affinity of this complex is relatively weak (K_d_ = 0.9 μM). Using Egl labeled with two different color Qdots, we showed that two Egls can bind to BicD in motile mRNPs at nanomolar concentrations, suggesting that interactions between components of the assembled complex likely enhance affinity. Because transported mRNAs in *Drosophila* extracts were shown to primarily contain a single mRNA (Amrute-Nayak and Bullock, 2012; Soundararajan and Bullock, 2014), Bullock and colleagues have proposed that a single mRNA may scaffold the association of two Egls with BicD CC3 to facilitate activation of dynein (McClintock et al., 2018). We showed that complexes with two Egl molecules had faster speeds and longer run lengths than mRNPs containing a mixture of 1 or 2 Egls. Separate experiments with dual-colored dyneins directly showed that the enhanced speed and run length can be attributed to the recruitment of two dyneins. Future stepping experiments will reveal if the molecular basis for the faster speeds is that the two-dynein complex exhibits fewer back and side steps, leading to a more efficient mini-ensemble. Our experiment showing that mRNPs with two Egls preferentially recruited two dyneins was striking because the overall stoichiometry used to form the core of the mRNP was one dynein per dynactin. The putative interaction between Egl and LC8 of dynein does not appear to contribute to dynein recruitment, because mRNPs reconstituted with dynein lacking LC8 were the same as those containing wild-type dynein. Our result is consistent with very recent studies showing that dynactin can recruit two dyneins to a dynein-dynactin-adaptor protein complex (Grotjahn et al., 2018; Urnavicius et al., 2018), and that complexes with two dyneins show faster movement and enhanced force (Urnavicius et al., 2018).

The cryo-EM studies of Grotjahn et. al. (Grotjahn et al., 2018) visualized two dimeric dyneins bound to essentially all (>97%) of dynein-dynactin-BicD2^CC1^ or dynein-dynactin-Hook3 complexes bound to a microtubule in the presence of AMP-PNP. Interestingly, Carter and colleagues (Urnavicius et al., 2018) also showed that BicDR1 and Hook3 recruited two dimeric dynein motors, but found that only 18% of the BicD2 complexes bound two dyneins, which they related to the positon of the N-terminus of BicD2. Our results showed the same number of two-dynein complexes with BicD versus Hook. Factors that influence recruitment of the second dynein to *Drosophila* BicD and human BicD2 remain to be determined. Our studies with full-length *Drosophila* BicD suggest that one factor influencing recruitment of two dimeric dynein motors in an mRNP is the presence of two bound Egls, thus establishing a link between cargo and number of bound motors. This *in vitro* reconstitution of an mRNP, from motor to bona fide biological cargo, provides a model system to test further interactions between components of the complex, which likely are stabilized by multiple weak interactions that synergize to produce a robust transport complex.

## Materials and methods

### DNA constructs

Full-length *Drosophila* BicD (NP_724056.1 or NM_165220.3) was cloned into pACSG2 for production of recombinant virus and expression in Sf9 cells. A FLAG tag at the C-terminus facilitated purification. Where indicated, constructs contained an N-terminal monomeric YFP, or a biotin tag for conjugation to a streptavidin-Qdot (Invitrogen). The biotin tag is a 88 amino acid fragment of the biotin carboxyl carrier protein (Cronan, 1990). Truncated human BicD2^CC1^ (NM_015250 and NP_056065.1), amino acids 25-398, was cloned into bacterial expression vector pET19 with either an N-terminal HIS and biotin tag, or an N-terminal HIS and monomeric YFP tag. Human BicD2^CC1^ aligns with *Mus musculus* BicD amino acids 25-400. Other *Drosophila* BicD truncation constructs were cloned with an N-terminal FLAG and biotin tag, or an N-terminal FLAG and monomeric YFP tag, and inserted into pACSG2 for recombinant virus production and Sf9 cell expression. BicD CC2 is amino acids L318-Q557, and BicD CC2-CC3 is amino acids L318-F782. Human Hook1 (NP_056972.1 & NM_015888.4) was cloned into pFastBac for recombinant virus production and expression in Sf9 cells with an N-terminal EGFP tag and a FLAG tag at the C-terminus. *Drosophila* Egalitarian (AAB49975.2 and U86404.2) was cloned into pACSG2 with either a C-terminal FLAG, or C-biotin followed by a HIS tag. The N-terminal 402 amino acids of mouse kinesin (NP_032475 and NM_008449.2) with a G235A point mutation was cloned into pET21b for expression of a rigor kinesin used for attachment of microtubules to the flow cell surface. Codon optimized human dynein for expression in Sf9 cells (DYNC1H1 (DHC), DYNC1I2 (DIC), DYNC1LI2 (DLIC), DYNLT1 (Tctex), DYNLRB1 (Robl) and DYNLL1(LC8)) was a generous gift from Simon Bullock (Schlager et al., 2014). The heavy chain was modified to contain an N-terminal FLAG tag followed by either a biotin or SNAP tag to enable labeling of the heavy chain. Separate recombinant viruses were produced to express each of the subunits (except for Robl and Tctex which were present in the same virus). All subunits were under the polyhedron promoter except for Robl which was under the p10 promoter.

### Reagents used for protein purification

Reagents used for protein expression and purification include: 4-(2-aminoethyl)benzenesulfonyl fluoride (AEBSF, Fisher BioReagents 30827-99-7), phenylmethylsulfonyl fluoride (PMSF, Sigma-Aldrich P7626), Tosyl-L-lysyl-chloromethane hydrochloride (TLCK, Sigma-Aldrich T7254), leupeptin (Thermo Scientific 78435), benzamidine (Sigma-Aldrich B6506), FLAG affinity resin (Sigma-Aldrich A2220), FLAG peptide (Sigma-Aldrich, F3290) and HIS-Select resin (Sigma-Aldrich P6611), and biotin (Sigma-Aldrich B4639).

### Protein expression and purification

Cytoplasmic dynein and dynactin were purified from bovine brain as described previously (Bingham et al., 1998), except that the preparation was scaled down to 1.5 brains (∼375 g). Alternatively, dynein was expressed in Sf9 cells as described below. Bovine tubulin was purified from brain tissue as described previously (Castoldi and Popov, 2003). Protein concentrations were determined using Bradford reagent with BSA as standard.

Dynein and accessory chains were co-expressed in Sf9 cells for ∼72 hours at 27°C, harvested, and re-suspended in 10 mM imidazole, pH 7.0, 0.3 M NaCl, 1 mM EGTA, 5 mM MgCl_2_, 7% sucrose, 2 mM DTT, 0.5 mM AEBSF, 0.5 mM PMSF, 0.5 mM TLCK, 5 μg/ml leupeptin, and 1.3 mg/ml benzamidine. Cells were lysed by sonication, and centrifuged at 257,000xg for 40 min. The clarified lysate was added to 4 ml FLAG affinity resin and incubated with mixing for 40 min. Resin was transferred to a column and washed with 200 ml FLAG wash buffer (25 mM imidazole, pH 7.4, 0.2 M NaCl, 1 mM EGTA) and eluted with the same buffer containing 0.1 mg/ml FLAG peptide. Peak fractions were concentrated using a Millipore Amicon Ultra-15 centrifugal filter and dialyzed against 5 mM NaP_i_, pH 7.2, 0.2 M NaCl, 1 mM DTT, 0.1 μg/ml leupeptin, and 50% glycerol for storage at -20°C.

BicD2^CC1^ with a N-terminal HIS and biotin tag was expressed in BL21(DE3) bacterial cells. Cells were induced with 0.7 mM IPTG and grown overnight at room temperature in LB broth containing 0.024 mg/ml biotin. Cells were harvested, pelleted, and re-suspended in HIS lysis buffer (10 mM NaPO_4_, pH 7.4, 0.3 M NaCl, 0.5% glycerol, 7% sucrose, 7 mM β-ME, 0.5 mM AEBSF, and 5 μg/ml leupeptin). Cells were lysed by sonication, clarified at 33,000xg for 30 min, and the supernatant bound to 3.5 ml of HIS-Select column. The column was washed with wash buffer (10 mM NaPO_4_, pH 7.4, 0.3 M NaCl) containing 10 mM imidazole, followed by 4 column volumes of wash buffer containing 30 mM imidazole. Protein was eluted in wash buffer containing 200 mM imidazole, and concentrated using a Millipore Amicon Ultra-15 centrifugal filter. Purified protein was clarified 487,000xg for 20 min and dialyzed against 25 mM imidazole, pH 7.4, 300 mM NaCl, 1 mM EGTA, 50% glycerol, 1 mM DTT, 0.1 μg/ml leupeptin for storage at -20°C.

Full-length *Drosophila* BicD was expressed in Sf9 cells. For constructs with a biotin tag, the media was supplemented with 0.2 mg/ml biotin. Cells were grown for ∼72 hours at 27°C, pelleted, and re-suspended in 10 mM imidazole, pH 7.4, 0.3 M NaCl, 1 mM EGTA, 2 mM DTT, 0.5 mM AEBSF, 0.5 mM PMSF, 0.5 mM TLCK, 5 μg/ml leupeptin, 1.3 mg/ml benzamidine. Cells were lysed by sonication and centrifuged at 257,000xg for 40 min. The clarified lysate was added to the FLAG affinity resin, and incubated with shaking at 4°C for 40 min. The resin was transferred to a column and washed with 200 ml FLAG wash buffer (25 mM imidazole, pH 7.4, 0.2 M NaCl, 1 mM EGTA) and eluted with FLAG wash buffer containing 0.1 mg/ml of FLAG peptide. Peak fractions were concentrated using a Millipore Amicon Ultra-15 centrifugal filter and dialyzed against storage buffer (10 mM Imidazole, pH 7.4, 200 mM NaCl, 1 mM EGTA, 50% glycerol, 1 mM DTT, 0.1 μg/ml leupeptin) for storage at -20°C. Hook1 was expressed and purified as described for BicD.

Egl containing a C-terminal biotin and FLAG tag, or only a FLAG tag, was expressed in Sf9 cells and purified similarly except that cells were infected for only 48 hours. Lysis and wash steps were done in buffers containing 1 M NaCl, and the protein was stored by snap freezing in liquid nitrogen and stored at -80°C.

For Egl-BicD co-expression in Sf9 cells, Egl contained a C-terminal biotin and HIS tag, and BicD contained N-terminal YFP and FLAG tag, or only an N-terminal FLAG tag. Following infection, cells were grown for ∼72 hours at 27°C and then harvested. The pellet was re-suspended in HIS lysis buffer (10 mM NaPO_4_, pH 7.4, 0.25 M NaCl, 0.5% glycerol, 7% sucrose, 0.1% NP40, 0.5 mM DTT, 0.5 mM AEBSF, 0.5 mM PMSF, 0.5 mM TLCK, 5 μg/ml leupeptin, 1.3 mg/ml benzamidine. The cells were lysed by sonication and the lysate was centrifuged 257,000xg for 40 min. A 5 ml HIS-Select column was prepared by washing the resin with 5 column volumes of water, 3 column volumes of 0.5 M imidazole pH 7.4, 20 column volumes of water, and re-equilibrated in 5 column volumes of HIS lysis buffer without DTT and protease inhibitors. The high imidazole wash allows for subsequent use of DTT. The clarified lysate was incubated with resin for 40 min and then washed in a column with ∼100 ml of 10 mM imidazole wash buffer (10 mM NaPO_4_, pH 7.4, 0.25 M NaCl, 10 mM imidazole, 0.5 mM DTT). Protein was eluted in 10 mM NaPO_4_, pH 7.4, 0.3 M NaCl, 200 mM Imidazole, 0.5 mM DTT and dialyzed overnight against 10 mM imidazole, pH 7.4, 0.25 M NaCl, 0.5% glycerol, 0.1% NP40, 1 mM DTT, 0.1 μg/ml leupeptin. The dialyzed protein was then purified over a FLAG column to remove excess Egl by incubating with FLAG affinity resin for 60 min, followed by washing and elution as described for BicD purification. Peak elution fractions were combined, dialyzed against 30 mM HEPES, pH 7.4, 0.25 M NaOAc, 2 mM MgOAc, 1 mM EGTA, 0.1 μg/ml leupeptin, and 2 mM DTT, snap frozen in liquid nitrogen, and stored at -80°C. Some co-expressed preparations were purified only on HIS resin with extensive washing to remove extra BicD. Elution fractions which showed approximately equal band intensities for BicD and for Egl were pooled and treated as described for the two column preparation. For negative stain electron microscopy, protein was dialyzed against 30 mM HEPES pH 7.2, 0.25 M KOAc, 2 mM MgOAc, 1 mM EGTA, 1 mM TCEP, 0.1 μg/ml leupeptin, centrifuged 487,000xg for 20 min, and drop frozen into liquid nitrogen.

Rigor kinesin (G235A), a mutant that binds to microtubules but does not dissociate in the presence of ATP or support microtubule motility, was cloned into pET21a. Expression was induced with 0.4 mM IPTG overnight at room temperature in *E. coli* Rosetta (DE3) (Novagen) in Terrific Broth Media (Invitrogen 22711-022) containing kanamycin. Cells were harvested, re-suspended in lysis buffer (10 mM Hepes pH 7.5, 10 mM NaCl, 1 mM EGTA, 0.25 mM DTT with 0.5 mM AEBSF, 0.5 mM TLCK, and 5 μg/ml leupeptin), and lysed by sonication. After clarification, the buffer was adjusted to a final concentration of 0.2 M NaCl and loaded onto a HIS-Select column equilibrated with lysis buffer containing 0.2 M NaCl. The column was washed with the same buffer containing 10 mM imidazole, then eluted with lysis buffer containing 0.2 M imidazole. Peak fractions were dialyzed against storage buffer (50% glycerol, 10 mM imidazole, 0.2 M NaCl, 1 mM EGTA, 1 mM DTT, and 5 μg/ml leupeptin) for storage at -20°C.

### *K10* mRNA constructs and synthesis

The 3’UTR of *Drosophila K10* mRNA (NM_058143.3),105-1165 nucleotides past the stop codon, was cloned after the SP6 promoter in the pSP72 vector (Promega) followed by a poly16A tail and an EcoRV site to allow the vector to be bluntly opened for use as a template for RNA transcription. The 43 nucleotide transport/localization signal (TLS) zip code starts 679 base after the start of the 3’UTR (Serano and Cohen, 1995). *K10* mRNA constructs contain 574 bases before and 443 bases after the TLS. For the *K10* no zip construct, the TLS element (CTTGATTGTATTTTTAAATTAATTCTTAAAAACTACAAATTAA) was removed. A minimal *K10* mRNA construct (*K10*_min_) consists of 195 nucleotides that center the TLS element. A minimal *K10* mRNA construct lacking the zip code (*K10*_min_ no zip) is the same sequence without the TLS and contains an additional 43 bases of 3’UTR sequence immediately following *K10*_min_ so that *K10*_min_ and *K10*_min_ no zip are the same size. The DNA template was bluntly linearized and transcribed using a phage SP6 RNA polymerase (RiboMAX system, Promega). Labeling of the *K10* RNA was achieved by adding a mixture of Alexa Fluor 488 labeled UTP (Molecular Probes, Invitrogen) in a 1:10 molar ratio to unlabeled nucleotides. For the dual-colored mRNA experiment, a second aliquot of *K10* mRNA was labeled with Andy Fluor 647 labeled UTP (GeneCopoeia).

### Flow cell preparation

PEGylated coverslips were made using methods adapted from (Gestaut et al., 2008). Glass cover slides (Fisher Scientific 12-545-M) were plasma cleaned for 5 min and transferred to glass Coplin jars containing 1 M KOH and then placed in a sonicating water bath for 20 min. Slides were rinsed thoroughly with nanopure water, then 95% ethanol and dried using a nitrogen stream. Slides were then placed in glass Coplin jars containing 1.73% 2-methoxy(polyethyleneoxy)propyltrimethoxysilane (Gelest, Inc SIM6492.7 – 25g) and 0.62% n-butylamine (Acros Organics 109-73-9) in anhydrous toluene (Sigma-Aldrich 244511), prepared with glass pipettes. Coplin jars containing slides were then placed in plastic bags, purged with nitrogen and incubated for 1.5 hours at room temperature. Following this incubation, the slides were dipped successively in two beakers containing anhydrous toluene and dried using a nitrogen stream. The slides were immediately made into flow chambers, placed in 50ml tubes and stored at -20°C. This procedure produces a PEGylated slide surface that contains small gaps for the purpose of microtubule attachment.

### Single molecule Total Internal Reflection Fluorescence (TIRF) microscopy assays

Bovine tubulin was thawed and centrifuged at 400,000 × g for 5 min at 2°C. Tubulin concentration was determined using Bradford reagent and diluted to 100 μM in ice cold BRB80 (80 mM PIPES, pH 6.9, 1 mM EGTA, and 1 mM MgCl_2_) and supplemented with 1 mM GTP. For generating labeled microtubules, unlabeled tubulin was mixed with 1 μM rhodamine-labeled tubulin (Cytoskeleton, Denver, CO) for a final labeled/unlabeled ratio of 1:100. The tubulin mixture was polymerized by transferring to 37°C water bath for 20 min and stabilized by adding 10 μM paclitaxel (Cytoskeleton, Denver, CO). Stabilized microtubules were kept at room temperature for experiments performed that day. Microtubules could be stored at 4°C for use in experiments within 3 days.

Labeled or unlabeled microtubules were adhered to PEGylated flow chambers using rigor kinesin for attachment. Rigor kinesin was diluted to 0.2 mg/ml in buffer B (30 mM HEPES, pH 7.4, 25 mM KOAc, 2 mM MgOAc, 1 mM EGTA, 10% glycerol, 10 mM DTT) and added to PEGylated flow chambers for 10 min at room temperature. Flow chambers were then washed three times in buffer A and blocked with buffer A containing 2 mg/ml BSA, 0.5 mg/ml ?-casein and 0.5% pluronic F68. Paclitaxel stabilized microtubules were diluted 100-fold to a final concentration of in buffer B containing 10 μM paclitaxel, and added to flow chambers and incubated for 10 min at room temperature. Flow chambers were washed three times with buffer B containing 10 μM paclitaxel to remove unbound microtubules.

For DDB^CC1^ single molecule motility, BicD2^CC1^ containing an N-terminal biotin tag was diluted in buffer B300 (30 mM HEPES pH 7.4, 300 mM KOAc, 2 mM MgOAc, 1 mM EGTA, 0.5% pluronic F68, 20 mM DTT) and centrifuged 400,000 × g for 20 min. Protein concentration was determined using Bradford reagent and diluted to 1 μM in B300. Dynein, dynactin and BicD2^CC1^ were mixed to a final concentration of 100 nM in buffer Go50 (30 mM HEPES pH 7.4, 50 mM KOAc, 2 mM MgOAc, 1 mM EGTA, 2 mM MgATP, 20 mM DTT, 8 mg/ml BSA, 0.5 mg/ml ?-casein, 0.5% pluronic F68, 10 μM paclitaxel and an oxygen scavenger system (5.8 mg/ml glucose, 0.045 mg/ml catalase, and 0.067 mg/ml glucose oxidase; Sigma-Aldrich-Aldrich)). Streptavidin-conjugated 655 quantum dots (Invitrogen) were added at 200 nM and incubated with proteins on ice for 30 min. Samples were diluted in buffer Go50 to a final dynein concentration of 1 nM and added to microtubule adsorbed flow chambers for imaging.

For imaging complexes where *K10* mRNA is labeled, BicD was diluted in B300 and centrifuged 400,000 x g for 20 min. Egl was diluted in B300 supplemented with 40 mM DTT and incubated on ice for 1 hour before centrifuging 400,000 × g for 20 min. Alternatively, co-expressed BicD-Egl complexes were used. Protein was determined using Bradford reagent. BicD and Egl were combined at 1 μM in B300. 50 nM dynein and dynactin, 100 nM BicD-Egl and 50 nM *K10* mRNA, synthesized with an Alexa Fluor 488 UTP for visualization, was mixed in buffer Go150 (buffer Go50 adjusted to a final concentration of 150 mM KOAc) containing 10 units of RNase Inhibitor (Promega N261B) and 0.25 mg/ml tRNA from *E. coli* (Sigma-Aldrich R1753). The order of mixing is BicD-Egl, dynein-dynactin, RNase Inhibitor, tRNA, *K10* mRNA. The mixture was incubated on ice for 45 min and diluted to a final dynein concentration of 1 nM in Go80 (buffer Go50 adjusted to a final concentration of 80 mM KOAc) before imaging.

For imaging of complexes where the adapters are labeled, either BicD or Egl containing a biotin tag were used for conjugation to quantum dots for visualization. Mixtures contained 50 nM dynein and dynactin, 100 nM BicD and Egl, 50 nM unlabeled *K10* mRNA and 200 nM Streptavidin-conjugated 655 quantum dots (Invitrogen). Complexes were incubated on ice for 45 min and diluted to a final dynein concentration of 1 nM in Go80 before imaging.

For single molecule pull-downs showing co-localization of YFP-BicD and Egl on microtubules, BicD fused to an N-terminal YFP tag and Egl containing a C-terminal biotin tag were prepared as described above and pre-mixed at 1 μM in B300. Mixtures containing 50 nM dynein and dynactin, 100 nM BicD-Egl, 50 nM unlabeled *K10* mRNA and 200 nM streptavidin-conjugated 655 quantum dots (Invitrogen) were diluted in Go150 supplemented with 10 units of RNase Inhibitor (Promega N261B) and 0.25 mg/ml tRNA (Sigma-Aldrich R1753). Mixtures were diluted so that the final concentration of dynein is 1 nM and imaged. YFP was used to visualize YFP-BicD and 655 Qdots were used to visualize Egl on rhodamine-labeled microtubules.

For the dual-color Egl experiment, 1 μM of biotinylated Egl was mixed with either 1 μM streptavidin 565 nm or 655 nm Qdots for 10 min, blocked with a 10-fold molar excess of biotin, and then combined. The combined labeled Egl was then added to a dynein-dynactin-BicD mixture followed by *K10* mRNA. The final molar ratio of components is dynein:dynactin:BicD:Egl:K10 = 1:1:1:2:1. Mixtures were diluted in buffer Go50 so that the final concentration of dynein in the assay was 1 nM. The 565 nm and 655 nm channels were simultaneously recorded and combined later in ImageJ 1.47v.

For dual-color dynein motility experiments, expressed dynein with an N-terminal SNAP tag on the heavy chain was biotinylated with SNAP-biotin (New England BioLabs, S9110S). 2 μM SNAP-dynein was incubated with 4 μM SNAP-biotin substrate (5 mM sodium phosphate, pH 7.5, 140 mM NaCl, 1 mM DTT) for 30 min at 37^°^C. Excess reagent was removed by overnight dialysis at 4^°^C in 30 mM HEPES, pH7.4, 300 mM KOAC, 20 mM DTT, followed by clarification at 350,000xg for 20 min. Dynein was labeled with either 525 nm or 655 nm streptavidin Qdots (molar ratio of 1:2) for 20 min on ice, then blocked with a 20-fold molar excess of biotin for 10 min. For single molecule AMP-PNP pulldowns on microtubules to determine percent dual-labeled complexes, 1 μM dynein-biotin was labeled with 2 μM streptavidin Alexa Fluor 488 or Alexa Fluor 647 for 30 min on ice, followed by addition of a 20-fold molar excess of biotin to prevent further binding (Invitrogen). Equimolar amounts of the two different colored dyneins (200 nM total) were incubated on ice with dynactin at a molar ratio of 2:1. Then BicD, Egl and *K10* mRNA were added to the dynein-dynactin complex and incubated another 45 min. The final molar ratio of dynein:dynactin:BicD:Egl:*K10* mRNA in DDBE plus K10 mRNA complex was 2:1:1:2:2. RNase Inhibitor (Promega N261B) and tRNA from E. coli (Sigma-Aldrich R1753) were added. A minimal DDB^CC1^ complex was formed with similar stoichiometry (2:1:1). To observe movement, the complex was diluted in buffer Go50 to a final dynein concentration of 0.5-1 nM. Motion was observed on rhodamine-labeled microtubules using TIRF microscopy and images of the Qdots (565 nm and 655 nm) were recorded simultaneously. For single molecule pulldowns to show dynein stoichiometry, dynein was observed on unlabeled microtubules in the presence of 6 mM AMP-PNP.

### Imaging and data analysis

Imaging was carried out on a Nikon ECLIPSE Ti microscope, run by the Nikon NIS Elements software package, and equipped with through-objective type TIRF. The samples were excited with the TIRF field of 405/488/561/640 nm laser lines, and emission was split by an image splitter (561nm or 638nm dichroic) and recorded on two Andor EMCCD cameras (Andor Technology USA, South Windsor, CT) simultaneously at 5 frames/second with automatic focus correction. The final resolution is 0.1066 μm/pixel. Motile mRNPs were tracked with labeled mRNA or with adapters labeled with a Qdot, and run lengths were measured with ImageJ and the particle-tracking plug-in MTrackJ (Meijering et al., 2012). For all processivity assays, frequencies were generated by counting the total number of runs in movies acquired no more than 15 min after dilution of the mRNP mixture. The total number of runs was divided by the total microtubule length, time and final dynein concentration to generate a run frequency. Speeds were measured by tracking run trajectories every 0.2 sec with ImageJ using the particle-tracking plug-in SpotTracker (Sage et al., 2005).

### Negative stain electron microscopy and image processing

YFP-BicD or YFP-BicD-Egl (expressed and purified as intact complex) was imaged in by diluting to 10-25 nM in buffer containing 30 mM HEPES, pH 7.2, 250 mM KOAc, 2 mM MgOAc,1 mM EGTA, 1 mM TCEP. For experiments with mRNA, YFP-BicD-Egl complex was mixed with a 2 fold molar excess prior to dilution. A 3 μl volume of diluted samples were applied to UV-treated, carbon-coated copper grids and stained with 1% uranyl acetate. Micrographs were recorded using an AMT XR-60 CCD camera at room temperature on a JEOL 1200EX II microscope at a nominal magnification of 40,000. Catalase crystals were used as a size calibration standard. 2D image processing was performed using SPIDER software as described previously (Burgess et al., 2004).The global image average and variance of BicD shown in Figure 2C results from a reference free alignment of 4205 images. To compare BicD with BicD-Egl complex, 4205 images of BicD alone and 4117 images of BicD-Egl were combined into a single stack and subjected to reference free alignment. Aligned images were classified into 20 classes using K-means classification and classes showing the most common b-type appearance were selected and a substack of images containing only b-type images was generated. This stack (4955 images) was realigned and the BicD-EGl images from the stack (n=2484) were averaged (Figure 4B). Subtracting the equivalent average of BicD only images (n=2471) resulted in the image shown in Figure 4C. The heatmap shown in Figure 4C was created by marking the position of the Egl in 342 images of the aligned BicD-Egl stack described above. To generate the low resolution 3D map shown in Video 1, 200 2D class averages of BicD were aligned to a starting model consisting of a 2nd order Gaussian ellipsoid. The resulting model was then refined against 4205 raw images of BicD. The median filtered volume and PDB files (PDB ID: 2Y0G, 5AFU, 1YT3) were arranged manually using UCSF Chimera to create the final movie.

### Abbreviations

BicD, full-length *Drosophila* Bicaudal D; BicD2^CC1^, coiled-coil 1 domain of mammalian Bicaudal D2; DDB^CC1^, dynein-dynactin-mammalian Bicaudal D2^CC1^; DDB, dynein-dynactin-BicD complex; DDBE, dynein-dynactin-BicD-Egl complex; Egl, the mRNA binding protein Egalitarian; *K10*_*min*_,195 nucleotides of *K10* that center the TLS element; Qdot, quantum dot; TLS, transport/localization sequence

## Acknowledgements

This work was funded by National Institutes of Health grant GM078097 to KMT, and American Heart Association Grant 12SDG11930002 to MYA. We thank the NHLBI Electron Microscopy Core Facility for the use of their microscopes.

## Author contributions

TES designed and performed single molecule research, analyzed data, wrote the paper; NB designed and performed electron microscopy experiments, analyzed data, wrote the paper; MYA designed and performed single molecule research, analyzed data, and contributed to writing the paper; CSB and EBK performed cloning, protein expression, purification and characterization, and mRNA synthesis; HL designed and performed single molecule research, analyzed data, and contributed to writing the paper; TAS provided expertise and insight on dynein-dynactin and contributed to writing the paper; KMT designed research and wrote the paper.

## Author ORCIDs

Kathleen Trybus 0000-0002-5583-8500

**Video 1. Low resolution 3D map of negative stain EM data (related to Figure 3).**

The video shows the size comparison of the apparent loop (EM volume depicted in gray mesh) with existing structures for parts of the YFP-BicD-Egl complex. The map has not been validated, and serves only as a guide. YFP molecules at the N-terminus of BicD are depicted in green (PDB ID: 2Y0G). The blue molecule is the exonuclease domain of RNAseD (PDB ID:1YT3), which serves as a proxy for the proposed exonuclease-like domain of Egl (Mach and Lehmann, 1997; Moser et al., 1997; Navarro et al., 2004). The white coiled-coil is the BicD2 N-terminal fragment from the structure of the dynein-dynactin-BicD2 complex (Urnavicius et al., 2015) (PDB ID:5AFU), fitted into the straighter part of the looped structure. In this interpretation, the remaining loop bulge must consist of CC2 and CC3. Given that CC2 is contiguous in sequence with the end of CC1, CC2 is likely to be the bottom part of the “b” structure, while CC3 binds to a region near the middle of CC1.

**Video 2. mRNPs with two dyneins move faster and longer (related to Figure 7).**

The DDBE plus *K10* mRNA complex containing two dimeric dyneins (yellow due to co-localization of a red and green Qdot) moved 12.2 μm in 20.4 seconds at a speed of 0.6 μm/s on a microtubule track (unlabeled). For comparison, a single dynein (red Qdot) moved a shorter distance (4.5 μm) at a slower speed (0.36 μm/s). The image was magnified 2-fold and played at 6x real time.

